# Cross-modality supervised image restoration enables nanoscale tracking of synaptic plasticity in living mice

**DOI:** 10.1101/2022.01.27.478042

**Authors:** YKT Xu, AR Graves, GI Coste, RL Huganir, DE Bergles, AS Charles, J Sulam

## Abstract

Synaptic plasticity encodes learning as changes in the strength of synapses, sub-micron structures that mediate communication between brain cells. Due to their small size and high density, synapses are extremely difficult to image *in vivo*, limiting our ability to directly relate synaptic plasticity with behavior. Here, we developed a combination of computational and biological methods to overcome these challenges. First, we trained a deep learning image restoration algorithm that combines the advantages of *ex vivo* super-resolution and *in vivo* imaging modalities to overcome limitations specific to each optical system. Applied to *in vivo* images from transgenic mice expressing fluorescently labeled synaptic proteins, this restoration algorithm super-resolved diffraction-limited synapses, enabling identification and logitudinal tracking of synaptic plasticity underlying behavior with unprecedented spatial resolution. More generally, our method demonstrates the capabilities of image enhancement to learn from *ex vivo* data and imaging techniques to improve *in vivo* imaging resolution.

## Introduction

Synaptic plasticity is a widely studied model of behavioral learning and memory encoding (Huganir and Nicoll, 2013; Malinow and Malenka, 2002; Nicoll, 2017) that directly links molecular changes at synapses—the sites of neurotransmission and communication between neurons—to changes in the strength of neural circuits. For instance, potentiation of excitatory synapses is observed during the learning of complex tasks, while synaptic degradation is observed in many neurological diseases, including cognitive and motor disorders (Henley and Wilkinson, 2016; Picconi et al., 2003; Volk et al., 2015). Unfortunately, it is extremely difficult to define how changes in synaptic strength within complex neural circuits manifest as learning in behaving animals. Indeed, investigation of synaptic plasticity in living animals has been limited by both the optical resolution of modern microscopy and a lack of biological tools to physiologically measure synaptic plasticity *in vivo* at scale. Developing new methods that overcome these technical constraints is thus vital to expand our understanding of how behavioral learning is encoded in real-time across the billions of synapses within the brain.

Super Ecliptic pHluorin (SEP), a pH-sensitive variant of GFP that fluoresces at neutral pH and is quenched at acidic pH (Miesenbock et al., 1998), is a useful tool to investigate protein trafficking in real time in living animals. For instance, by fusing SEP to the extracellular domain AMPA-type glutamate receptors (AMPARs), we can directly visualize trafficking and recycling of these proteins at the synaptic membrane. This visualization is possible as the acidity of the internal cellular environment quenches the fluorescence of non-functional internalized receptors, whereas the functional complement of receptors fluoresce at neutral pH at the cell surface. Observing SEP-labeled AMPAR fluorescence *in vivo* has recently been used to track changes in synaptic strength in living animals (Chen et al., 2021; Graves et al., 2021; Roth et al., 2020; Tan et al., 2020; Zhang et al., 2015). Using a transgenic approach that fluorescently labels all endogenous synaptic proteins in a manner that minimizes disruptions to native protein expression and does not impair physiological function (Graves et al., 2021). Thus while it is now theoretically possible to comprehensively visualize all synapses in living mice to create an intricate and dynamic map of synaptic plasticity across the entire brain, significant technical barriers remain to be overcome in analyzing such imaging data.

For one, visualizing endogenous SEP-tagged receptors at synapses in living animals presents a substantial challenge. Synapses are sub-micron-diameter structures that exist in high-density across the brain (Bock et al., 2011), making it challenging to resolve individual synaptic structures. Moreover, as the intensity of SEP fluorescence is correlated with synaptic strength, the distribution of synaptic brightness is broad, such that a proportion of synapses will be rather dim. Thus, to image SEP signals longitudinally *in vivo*, a fine balance must be achieved between imaging resolution, depth, speed, and laser power. For instance, while two-photon microscopy (2p) enables investigation of biological processes in living animals, as near-infrared excitation enables deeper tissue penetration, the resolution achievable in 2p imaging falls far behind that of single photon microscopy *in vitro* (Denk and Svoboda, 1997). In particular, axial resolution is especially impaired in 2p microscopy, as there is no pinhole to enable optical sectioning, and the long working distance required for *in vivo* imaging means that high-numerical-aperture objectives cannot be used. Conversely, adapting *in vitro* super-resolution microscopy elements, such as Airyscan detectors, to *in vivo* imaging is also difficult, as depth-dependent light scattering, movement artifacts, and tissue swelling force compromises between acquisition resolution, size, depth, and photobleaching. As such, the current state-of-the-art only allows for live imaging of molecular synaptic changes in *ex vivo* slice preparations, as *in vivo* imaging lacks the resolution to adequately explore neural tissue at the molecular level.

To overcome these biological limitations, we used machine learning to combine the advantages of both *in vitro* and *in vivo* imaging modalities. The application of convolutional neural networks (CNNs) in image restoration presents one promising avenue to selectively balance the benefits of different imaging modalities and alleviate the limitations of physical optical elements (LeCun et al., 2015; Nehme et al., 2018; Weigert et al., 2018). Unlike traditional restoration algorithms (Arigovindan et al., 2013; Richardson, 1972), deep learning models are well suited for image restoration as they can learn application-specific information from training data, thereby adapting to the high complexity of imaging in living animals. However, the necessity to learn data statistics from paired training image samples (registered high- and low-resolution images of the same tissue) creates a challenge in scenarios where optimal-resolution data is lacking, as is the case for *in vivo* imaging of nanoscale structures (Weigert et al., 2018). Identifying the exact same synapses *in vivo* and then again at high-resolution *in vitro* with perfect registration is extremely challenging and has only been achieved in small tissue volumes (Hirabayashi et al., 2018; Micheva et al., 2016).

In the present study, we circumvented this limitation by developing a novel approach that produced cross-modality paired low- and high-resolution training data from live brain slices to train a U-net-based restoration algorithm (Content-Aware Image Restoration, CARE; Weigert et al., 2018). At a high level, rather than using data from the same modality to train the model, we learn a signal model for the tissue properties as observed in alternate conditions using different imaging modalities. This approach super-resolved diffraction-limited, fluorescently labeled synapses *in vivo* by enhancing low-resolution 2p-acquired datasets, yielding Airyscan-like super-resolution image quality. Overall, a significant improvement in the axial resolution of restored images, as well as the size, number, and precision of synaptic detections was achieved in super-resolved volumes acquired from living mice undergoing behavioral learning. Moreover, computationally enhanced synapses were easier to track over time, leading to fewer association errors, facilitating the tracking of behaviorally relevant changes in synaptic plasticity *in vivo*. We show that this paired, live-slice supervised image restoration pipeline can be used to super-resolve synapses over time in live-imaging experiments. This is the first application that enables longitudinal visualization of synapses within regions of high synaptic density over weeks *in vivo*. By combining the advantages of multiple imaging modalities, deep learning, and transgenic labeling of endogenous proteins, this platform provides a means to study diffraction limited objects (e.g., axon terminals or dendritic spines) in living animals, enabling new experimental paradigms to explore how dynamic regulation of synaptic strength encodes learning and memory.

## Results

### Transgenic labeling enables visualization of individual excitatory synapses in vivo

AMPA-type glutamate receptors (AMPARs) mediate the majority of fast excitatory neurotransmission in the mammalian brain and their dynamic regulation is a central mechanism of synaptic plasticity. To visualize synapse dynamics in behaving animals, we used CRISPR-Cas9 DNA editing to create a novel transgenic mouse line with fluorescently labeled AMPARs. In homozygous *SEP-GluA2* mice, every endogenous GluA2 AMPAR subunit is fused with Super Ecliptic pHluorin (SEP). By coupling SEP to the extracellular N-terminus of the receptor, only GluA2-containing receptors at the cell surface are fluorescent, as fluorescence of internalized receptors in acidic compartments is quenched (**Fig. 1a**). Importantly, as an overwhelming majority of excitatory synapses contain GluA2 (Diering and Huganir, 2018), this transgenic line enables visualization of nearly all excitatory synapses in the brain. Imaging synapses using super-resolution confocal microscopy with Airyscan detectors in live-tissue slices in which some neurons expressed tdTomato confirmed that SEP-GluA2 puncta colocalize with dendritic spines *in vitro* (**Fig. 1b-d**), validating the spatial association of SEP fluorescence with physical synaptic sites. Further, using 2p microscopy through cranial windows *in vivo*, we visualized dense networks of millions of individual synapses in living mice (**Fig. 1e-f**).

**Figure 1.**
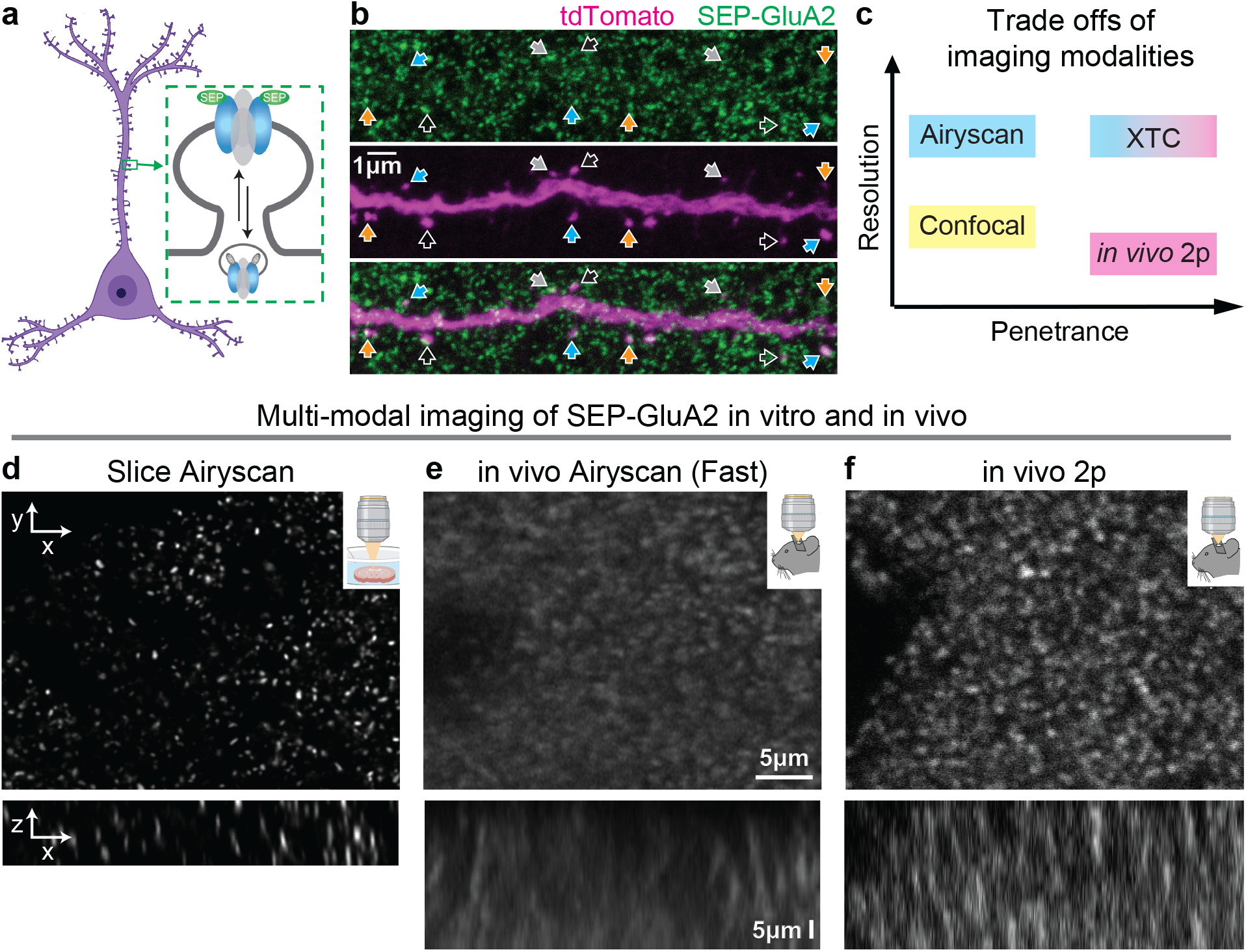
Resolving millions of individual synapses *in vivo* requires depth and resolution. (**a**) CRISPR-based transgenic labeling of the GluA2 AMPA receptor subunit with a pH-dependent fluorescent tag (Super Ecliptic pHluorin, SEP) enables *in vivo* visualization of endogenous GluA2-containing synapses, located in spines throughout the dendritic arbor. (**b**) Single high-resolution imaging plane from fixed slice tissue with endogenous fluorescence, acquired using Airyscan detectors. Dendrite expressing tdTomato (magenta) enables visualization of SEP-GluA2 puncta (green) colocalization with dendritic spines. Arrows mark examples of SEP-GluA2/spine overlap. (**c**) Tradeoffs of different imaging modalities. Airyscan allows high resolution, but with slow speed and limited depth. Conversely, *in vivo* 2p resonance imaging allows high scan speed and deeper tissue penetrance, but with lower resolution. XTC combines the advantages of different modalities. (**d - f**) Example XY projections (top) and XZ projections (bottom) of different imaging modalities. *In vivo* microscopy suffers from substantial blur due to extended working distance and low NA, compared to in vitro super-resolution modalities.

Labeling nearly all excitatory synapses theoretically enables a comprehensive analysis of synaptic plasticity underlying behavior, but represents a several-orders of magnitude increase in data complexity compared to recent studies using overexpression of SEP-AMPARs in a sparse subset of neurons (Chen et al., 2021; Roth et al., 2020; Tan et al., 2020; Zhang et al., 2015). Unfortunately, optically resolving this dense and relatively dim endogenous SEP signal in living animals is exceptionally challenging, particularly due to motion artifacts associated with *in vivo* imaging. We tested several imaging modalities (2p with galvanometric scanner, 2p with resonance scanner, Airyscan *in vivo*, Airyscan Fast *in vivo)* (**Fig. 1c–e**), but were unable to achieve sufficient resolution to reliably resolve adjacent synapses *in vivo*, particularly in the axial plane. While the best resolution *in vivo* was achieved using 2p excitation and a resonance scanner that reduced motion artifacts, none of these methods preserved the overall shape and clarity of fluorescent synapses observed using super-resolution Airyscan microscopy in brain slice preparations (**Fig. 1d**). Thus, to improve synapse detection in 2p imaging datasets, we sought to combine the resolution of Airyscan microscopy with the speed and penetration of 2p excitation using computational image restoration (**Fig. 1c**).

### Improved resolution of individual synapses using posthoc supervised CNN image restoration

Computational image restoration presents an efficient, highly adaptable avenue for overcoming the limitations associated with specific optical systems (LeCun et al., 2015; Nehme et al., 2018; Weigert et al., 2018). Using a supervised training paradigm, whereby a CNN is directed to enhance the image quality of a suboptimal dataset to that of a higher resolution target dataset, researchers can, in theory, selectively balance the advantages of different imaging modalities by mapping them together using paired high- and low-resolution images. In our case, we want to combine the speed and penetrance of 2p microscopy, which facilitates *in vivo* imaging, with the substantially higher resolution of Airyscan microscopy, which has poor penetrance and cannot be widely applied *in vivo*. Unfortunately, acquiring such a paired dataset is nearly impossible in our application, as the low-resolution 2p datasets represent the upper limit of data quality in current *in vivo* optical applications (**Fig. 1**). Thus, to sufficiently resolve millions of fluorescently labeled synapses *in vivo*, we turned to *in vitro* imaging of live, acute slices of SEP-GluA2 brains to produce a training dataset from which a restoration algorithm (Content-Aware Image Restoration, CARE; Weigert et al., 2018) could learn a mapping strategy from low-resolution single-photon confocal (*Slice 1p*) images to high-resolution Airyscan image quality (*Slice Airy;* **Fig. 2a**). We hypothesized that images from acute live tissue-slice preparations, acquired immediately after dissection in physiological buffers to preserve tissue quality, would be sufficiently similar to *in vivo* datasets such that a restoration CNN trained using paired live-slice training data could accurately enhance the resolution of in vivo images. Thus, we defined ground truth high-resolution data (*Slice Airy*) as images acquired with Airyscan super-resolution microscopy while low-resolution paired data (*Slice 1p*) was generated using 1p excitation with an open-pinhole, decreased laser power, and high gain, to resemble the axial blur and high noise of *in vivo* 2p imaging (**Fig. 2a-c**). A CNN with a modified UNet architecture (Ronneberger et al., 2015) was then trained using this paired high-low training data to generate a restoration model. Finally, after training the image restoration algorithm, we assembled a tracking pipeline to enable synapse tracking across longitudinal imaging experiments (**Fig 2d**).

**Figure 2.**
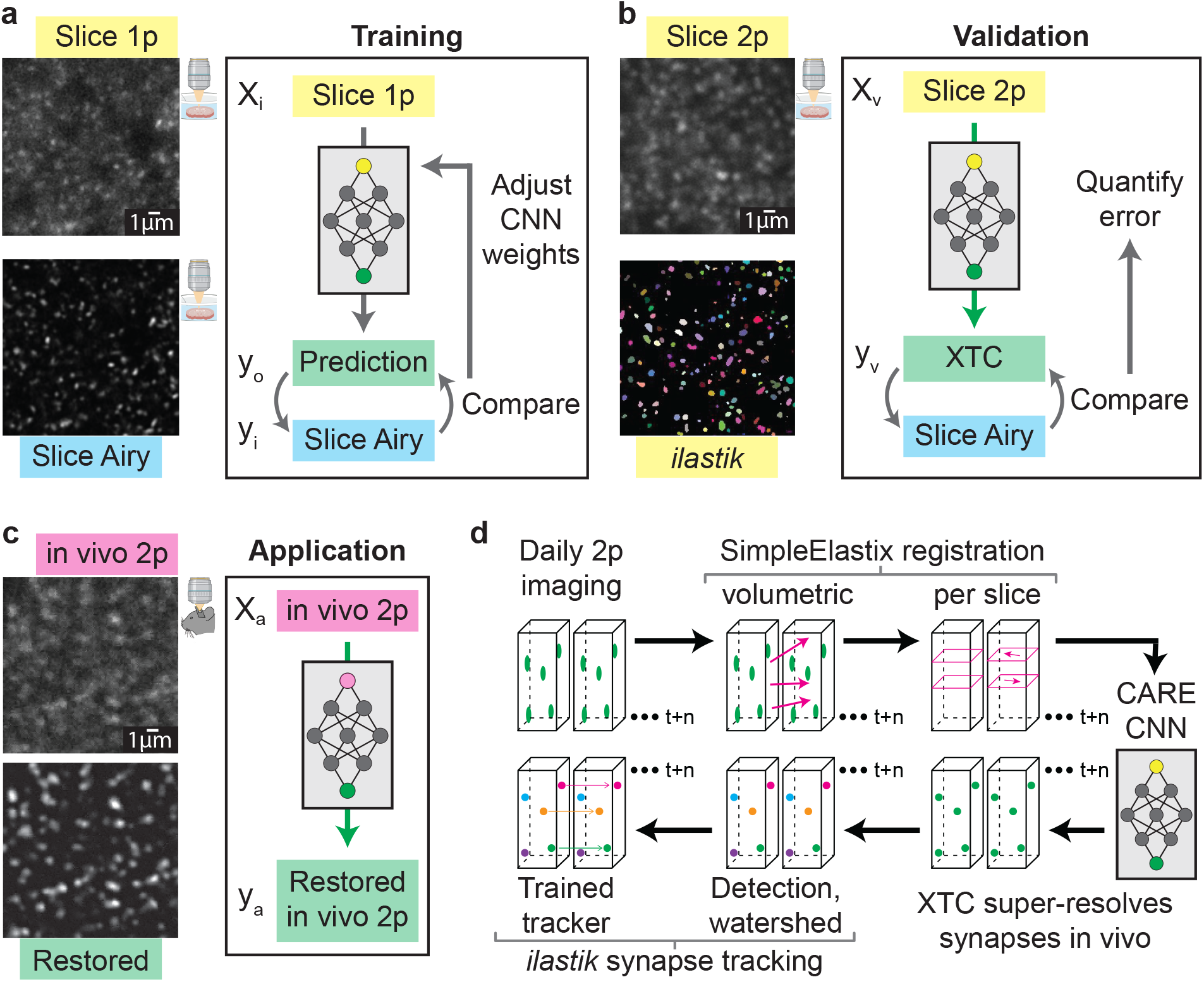
Cross-modality pipeline to train and implement XTC. (**a-c**) Diagrams of training, validation, and application workflow. Left, example images of each color-coded imaging modality. Right, workflow of training, validation, or application. (**a**) CNN was trained using 1p confocal images from acute slices of SEP-GluA2 tissue (x_i_). CNN output (y_o_) was compared to ground truth (high-resolution Airyscan imaging of the same tissue, y_i_) to improve network performance. (**b**). Network output was validated by comparing to ground truth and annotations by expert humans, enabling quantification of error rates. (**c**). Trained restoration CNN was applied to *in vivo* 2p images, restoring optimal “Airyscan-like” resolution to *in vivo* imaging volumes. (**d**) Pipeline for longitudinal tracking of fluorescently labeled SEP-GluA2 synapses *in vivo*. Daily imaging volumes were aligned using pairwise affine registration, followed by slice-by-slice pairwise affine registration to compensate for depth-dependent local tissue shift. Registered volumes were restored with the cross-modality trained XTC network. Millions of individual synapses were segmented with an *ilastik-trained* random forest model, followed by watershed to separate adjacent objects. Finally, a tracker trained through structured learning was used to longitudinally track synapses.

### Cross-modality image enhancement restores high-resolution synaptic signals

Preserving the intensity and size of SEP-labelled synapses is critical to provide an accurate assessment of AMPAR changes at these sites, as SEP fluorescence is directly correlated with synaptic strength (Graves et al., 2021). To assess how image restoration impacts the distribution of synapse shape, intensity, and spatial distribution throughout the brain, we generated validation data with paired high-resolution volumes, imaged with Airyscan microscopy (*Slice Airy)*, and low-resolution volumes, imaged with 2p excitation and resonance scanning (*Slice 2p*, **Fig 2a-b**).

*In vivo 2p* images restored using our multi-modal cross-trained CARE (*XTC*) model showed dramatic improvements in both lateral and axial resolution (Fig. 3a, Video 1). Moreover, enhanced volumes showed a similar spatial distribution of synapses compared to the low-resolution *in vivo 2p* data, indicating that the restored signal was not shifting the spatial distribution of synapses (Fig. 3a overlay). Closer inspection of distinct regions with sparse synapses, dense synapses, and near blood vessel occlusions or depth-dependent signal-loss, demonstrated that the CNN adapted to regional changes in image statistics to faithfully preserve the overall distribution of visible synapses (Fig. 3b).

**Figure 3.**
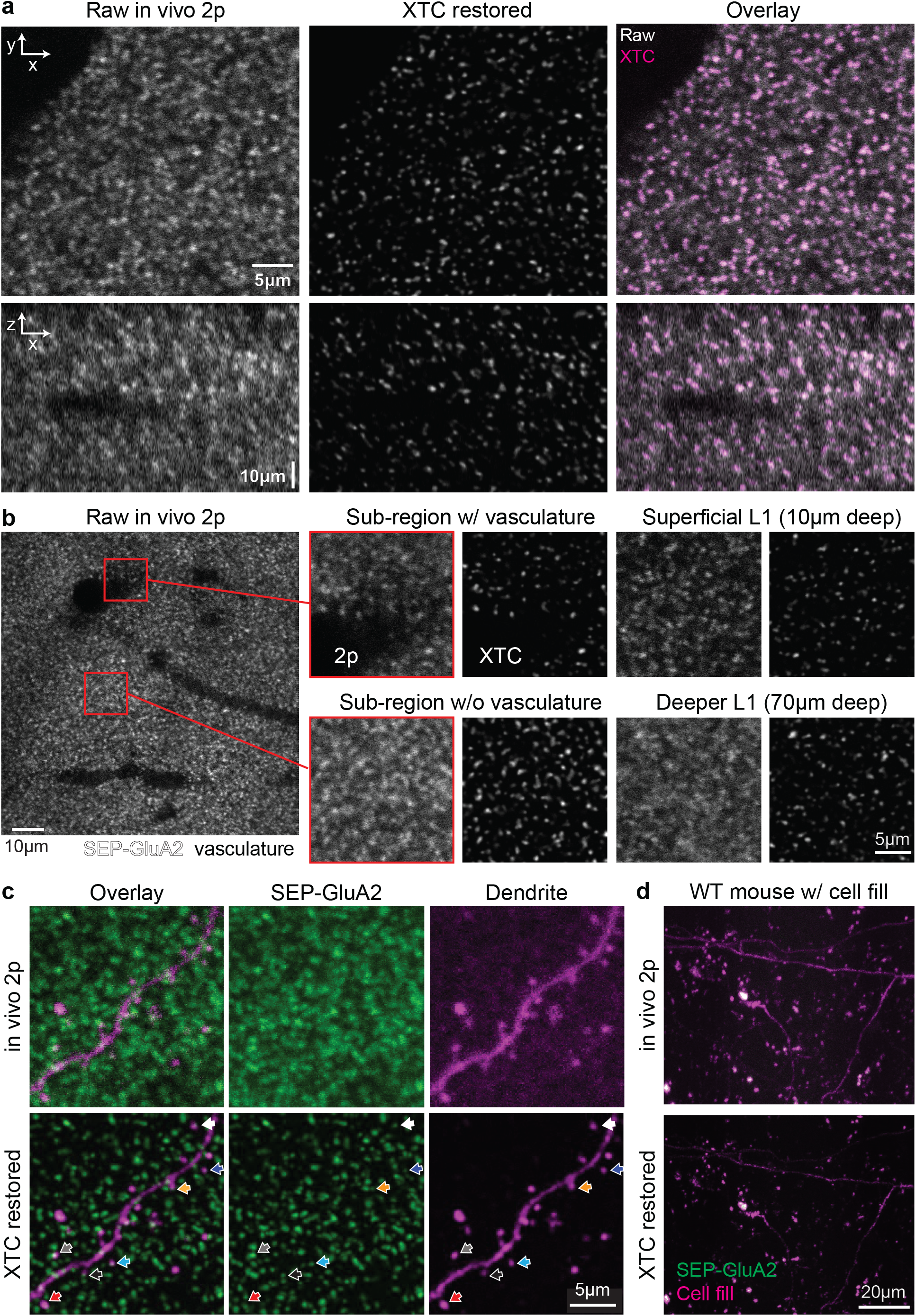
XTC super-resolves SEP synapses in vivo. (**a**) Comparison of same *in vivo* 2p image before (left) and after XTC (middle). Overlay of raw and XTC-processed data (right) shows high correspondence. (**b**) Representative slice from a single volume acquired *in vivo* (100 x 100 x 70 μm). Zoomed insets show XTC performance near blood vessel obstructions, in sparse and dense regions, and deeper in cortex. (**c**) SEP-trained XTC model generalizes to alternative imaging channels without creating artifacts. Super-resolved dendrite and associated spines are much easier to colocalize with XTC resolved SEP-GluA2 synapses (bottom, colored arrows show same synapse across image channels). (**d**) XTC does not detect signal in WT mice with unlabeled synapses, while preserving and enhancing filled dendrites (magenta), demonstrating the generalizability of XTC pipeline.

As the functions parametrized by CNNs are highly nonlinear and complex, it is important to empirically test for artifactual errors, such as “hallucinated” false-positive synapses not actually present in the data. We thus tested XTC on different fluorescence signals to determine if our model generalized well beyond the data distribution used for training. First, we applied XTC to a structural neuronal signal: viral expression of the fluorescent protein tdTomato to achieve a sparse cell fill. We observed that restored images faithfully retained the linear properties of dendrites, despite not having been trained on structural images containing dendrites, and did not create false-positive spherical “synapse-like” objects (Fig. 3c). The restoration model thus appeared to generalize well to this new neuronal fluorescence and was able to denoise both the dendrites and associated spine heads. Moreover, after enhancing both the tdTomato and SEP signals independently using XTC, SEP-labelled synapses were colocalized with dendritic spines showing that the spatial distribution of SEP synapses was consistent with their expected biological location after image restoration (Fig. 3c). Finally, we also tested our model on wild-type animals (not expressing SEP) with tdTomato-filled neurons. Again, we observed that the enhanced data did not create false-positives in the wild-type green SEP imaging channel (Fig. 3d). Overall, our tissue-slice trained image restoration algorithm faithfully enhanced synaptic fluorescent signals *in vivo* without distorting their underlying shape and visualized distribution.

To compare individual synapses before and after image enhancement to synapses from *Slice Airy* validation data, we trained voxel-wise random forest classifiers using *ilastik* (Berg et al., 2019) to volumetrically segment individual synapses. Three *ilastik* models were trained using sparse human annotations to detect synapses in *Slice 2p, XTC Restored*, and *Slice Airy* (ground truth) volumes, respectively (**Fig. 4a,b**). The low complexity of off-the-shelf *ilastik* classifiers, each trained with only 30 sparse human annotations, allowed us to assess how XTC restoration simplifies the challenge of synapse detection. For instance, given a trivial task, such as detecting synapses in optimal images with unlimited resolution and no noise, a simple binary classifier would be able to accurately segment all synapses.

**Figure 4.**
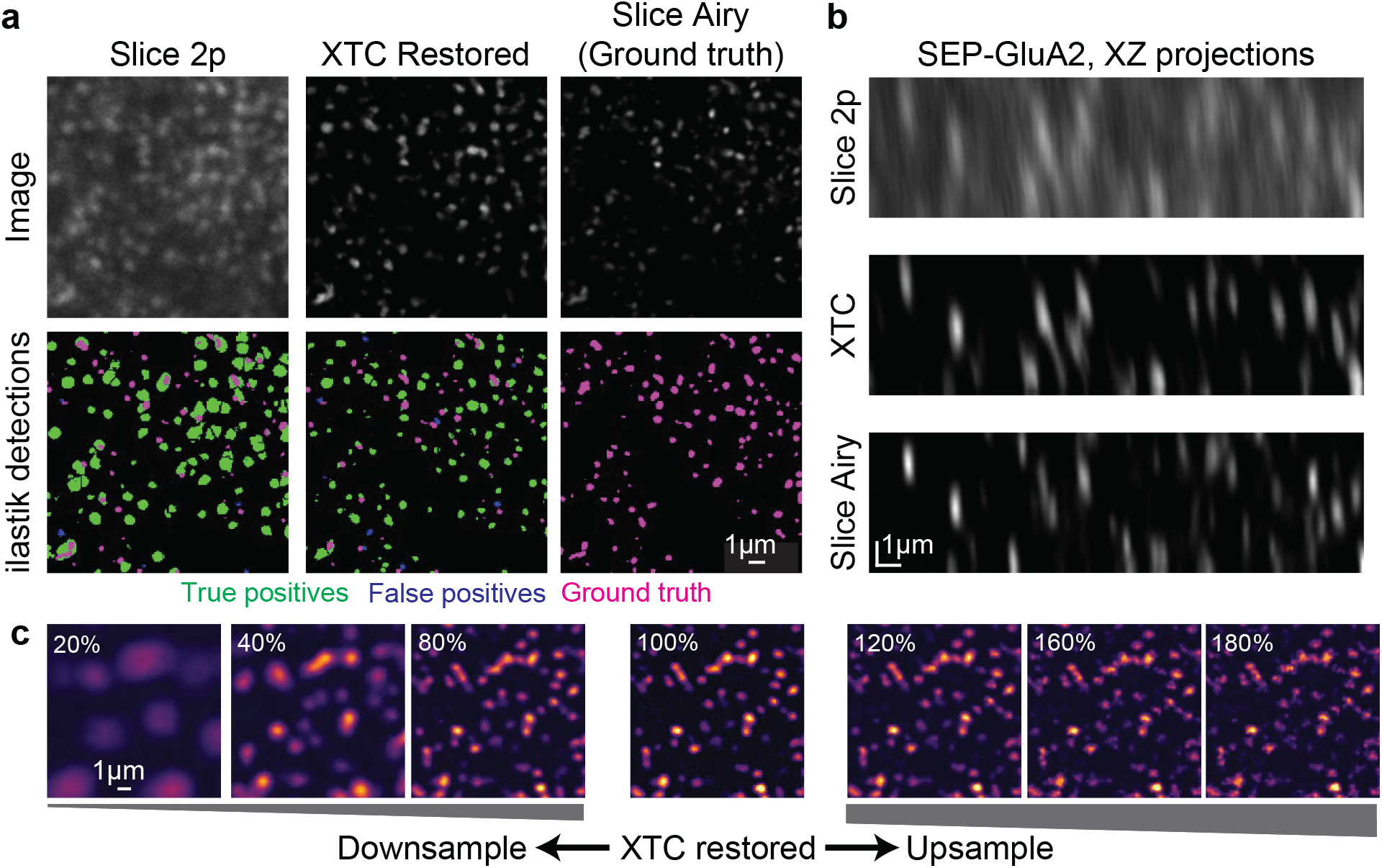
XTC enables high-precision segmentation of individual synapses. (**a**) Top, comparison of signals throughout XTC workflow used for validation. Bottom, synapse detections from top row volumes using trained *ilastik* synapse segmentation models. True positives (green), false positives (blue), and false negatives (magenta) are indicated. (**b**) XTC restoration improves lateral resolution as well. (**c**) Effects of up- and down-sampling the input 2p data (input) on XTC performance. Ideal resolution should be within range of training data resolution (0.063 - 0.096 μm/px in XY and 0.33 μm/px in Z).

However, accurate segmentation becomes more challenging in images with low resolution and high noise, and the performance of a model trained with the same number of human annotations deteriorates. Thus, when comparing the performance of our ilastik classifiers on each imaging dataset, we concluded that XTC restoration greatly simplifies the challenge of synapse segmentation, as each individually matched synapse, between *XTC Restored* and *Slice Airy*, was more similar in size, shape, and intensity than each matched synapse between *Slice 2p* and *Slice Airy* images (**Fig. 4a-b**). Finally, we also assessed the impact of varied image resolution on our model’s ability to generalize to datasets with different scales. We found that image enhancement performed best within the resolution range of the CNN training data (0.063 - 0.095 μm/px XY), indicating that the model should always be used within this range (**Fig. 4c**). At significantly lower resolutions (20% of original), we observed over-simplification of detections, and at higher resolutions (180% of original), we observed spotty artifacts that appeared as false synapses (**Fig. 4c**). Given training data at different resolutions, the CNN model would be expected to generalize to other use cases at different scales.

After initial visual inspections, we also quantitatively compared the intensity, size, and overlap of volumetric synapse segmentations before and after CNN restoration to ground-truth *Slice Airy* detections. First, we compared the mean intensity of each detection that matched with ground-truth in either *Slice 2p* or *XTC Restored* volumes. We found that image restoration preserved the correlation of mean intensity values for individual synapses relative to ground truth (**Fig. 5a,b**; r = 0.59 and 0.68, respectively). To compare these correlations to a semi-randomized baseline, we rotated the *Slice Airy* volume (*Rotated*) and found that the mean intensities of matched detections were no longer correlated between *Rotated* and unrotated *Slice Airy* volumes (**Fig. 5c**; r = −0.14). Next, we assessed the distribution of total sum intensity for individual synapses in each imaging condition and found that the distribution of XTC Restored synapses better matched the distribution of synapses in ground-truth data (**Fig. 5d,e**). Moreover, the overall shape of XTC Restored synapses was better matched to ground-truth detections than synapses from either *Slice 2p* or *Rotated* volumes, as indicated by the Jaccard overlap index (Jaccard, 1912) (**Fig. 5f**). When we examined the distribution of true positive (TP), false negative (FN), and false positive (FP) detections between XTC Restored and ground-truth volumes, we observed that the FP rate was extremely low. Although the FN rate was relatively high, most of these missed detections were either extremely small or dim (**Fig. 5g, h**). This suggests that image restoration cannot surpass the physical limits of optical elements and thus synapses that are very small or dim (< ~0.3 μm diameter) cannot be reliably detected *in vivo*, even after XTC restoration. Thus, this approach provides high-confidence detection of synapses with relatively higher AMPAR content. Finally, the structural similarity index (SSIM) was highest between XTC Restored and ground truth volumes, indicating that the overall structure and distribution of intensities is preserved with XTC processing (Fig. 5i). Overall, this quantitative assessment indicates that XTC processing (1) does not shift the relative intensity values underlying SEP signals, (2) improves the size and shape of synapse segmentations by simplifying the challenge of voxel classification, and (3) has a low FP rate and a high FN rate that is associated with the lower physical limit of *in vivo* imaging.

**Figure 5.**
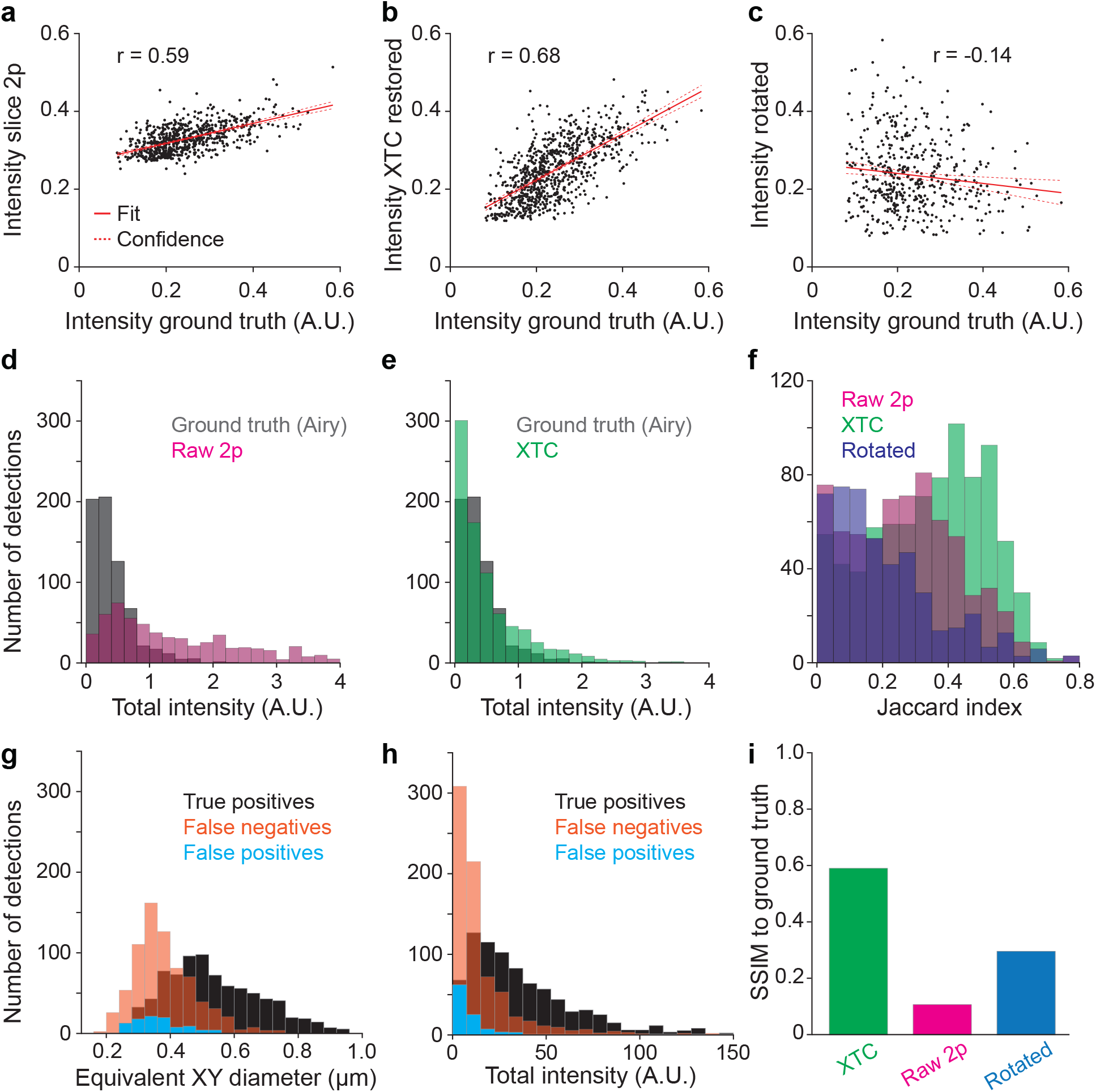
Validation of automated synapse detection following XTC. (**a-c**) Comparison of mean fluorescence intensity of matched synapses from *Slice 2p, XTC Restored*, and *rotated* volumes to the ground truth volume supports that restoration does not alter intensity values. (**d,e**) *XTC Restored* synapses better match ground truth detections for total intensity (sum of total voxel intensity within segmentation) than detections made in *Slice 2p*. (**f**) Jaccard index overlap of paired detections also supports that *XTC Restored* detections display high overlap with ground truth detections. The distribution of synapse diameter (**g**) and total intensity (**h**) for paired true positive, false negative, and false positive detections in *XTC Restored* volume relative to ground truth synapses detected in *Slice Airy* validation data. XTC restoration showed high precision (low false positive rate). (**i**) SSIM with ground truth data was greatly increased with XTC restoration.

#### Cross modal image enhancement facilitates longitudinal tracking of synaptic plasticity

Accurately tracking millions of synapses over extended periods of time in behaving animals is an important goal that may help define changes in synaptic plasticity responsible for learning and memory. Thus, we assessed if labeled synapses could be registered and tracked over weeks of imaging. We hypothesized that XTC image restoration would greatly facilitate our ability to track individual synapses by improving SNR and reducing tracking ambiguities. To perform a comparison of synapse tracking before and after image enhancement, a general pipeline for object detection and association was established (Fig. 2d and 6a). Cranial windows were surgically implanted over retrosplenial cortex and animals were imaged every 1-2 days for nearly two weeks to longitudinally track changes in synaptic plasticity at baseline and during behavior. Images were then registered to each other volumetrically, followed by a slice-by-slice basis using affine transformations within the *SimpleElastix* python package. After registration, only minor shifts in tissue movement persisted (Fig. 6c). We then masked out portions of the image volume that were obscured by blood vessels to prevent ambiguous synapse associations. The volumes were then restored with XTC, and synapses were volumetrically segmented using a trained *ilastik* classifier and subsequent watershed segmentation. Finally, to determine the association between detected synapses across multiple days, we trained a synapse tracking algorithm in *ilastik* using structured learning with sparse human annotations (Berg et al., 2019) (Fig. 6b - c).

**Figure 6.**
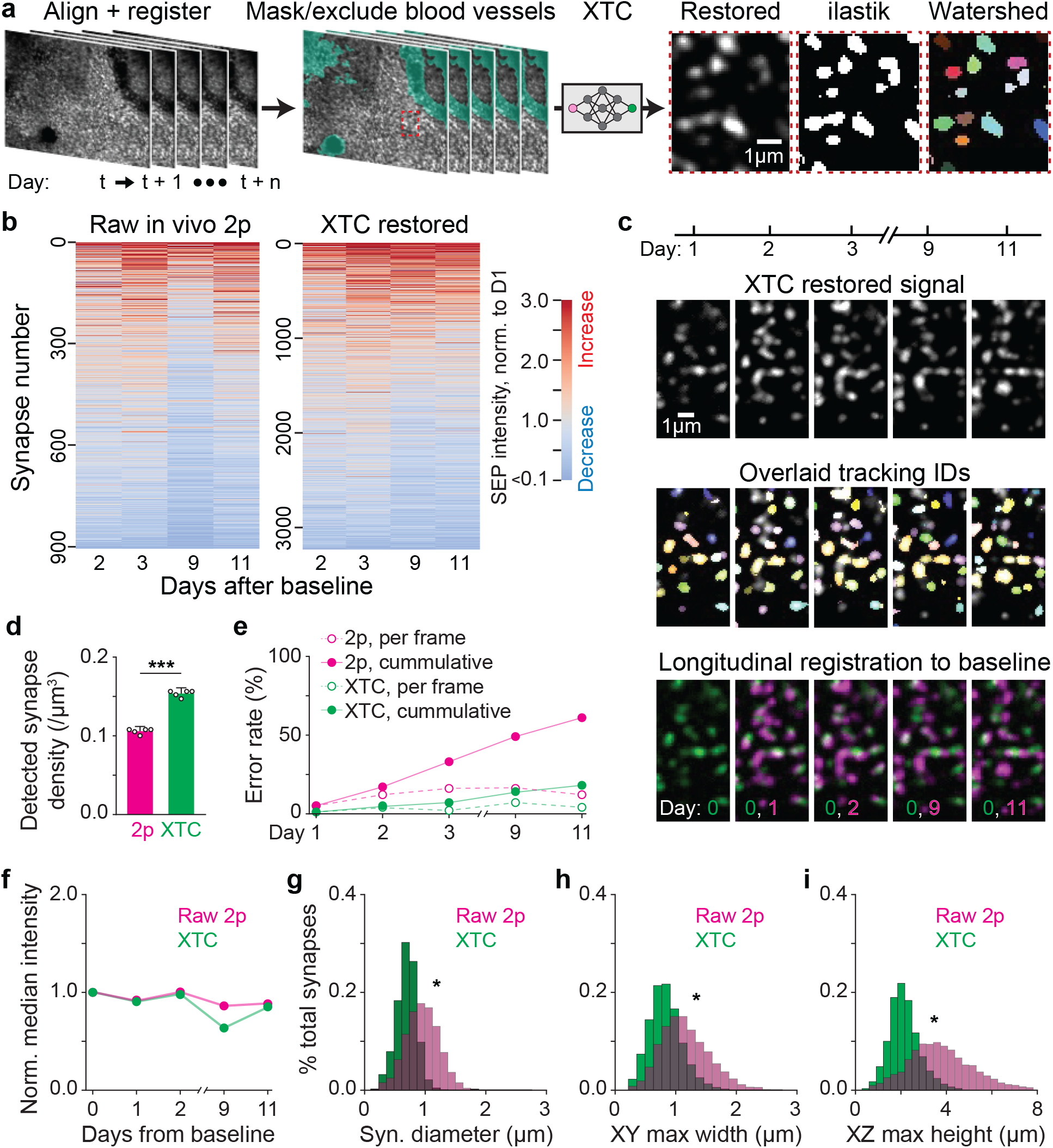
XTC enables tracking of thousands of registered synapses across days during behavior. (**a**) Overview of preprocessing pipeline for tracking. Volumes were acquired using *in vivo* 2p microscopy over 10 days and registered to each other. Blood vessels were masked to facilitate tracking. Imaging volumes were super-resolved with XTC; synapses were detected and separated with watershed segmentation. (**b**) Relative fold change of individual tracked synapses before and after XTC processing across 10 days of imaging. Fold change relative to Day 0 baseline. (**c**) Examples of single-synapse-resolution registration and tracking *in vivo*. Top, representative raw signal across 10 imaging days. Middle, same fields of view, with synapse detections overlaid. Identical colors across different days indicates the same tracked synapse. Bottom, registered and aligned signals (magenta), relative to Day 0 (green). (**d**) The overall density of synapse detections at each timepoint was increased after XTC restoration. (**e**) Error rate of tracking for 100 randomly selected synapses, as curated by expert humans, from *in vivo* 2p and XTC processed volumes. (**f**) Median fold change of all synapses at each timepoint was steady for XTC Restored data but showed a large dip on day 9 for *in vivo* 2p data suggesting tracking errors. (**g–i**) Distribution of diameter in XY, major axis length in XY, and major axis length in Z for individual synapses in both *in vivo* 2p and XTC Restored volumes.

Synapse tracking was greatly improved when XTC was applied to super-resolve *in vivo 2p* volumes. The number of synapses tracked over 10 days was more than 3-fold higher in *XTC Restored* volumes (Fig 6b; 907 vs. 3241 detections) and the overall density of accurate synapse detections at each timepoint was also increased by 50% (Fig. 6d; 0.106 ± 0.003 synapses/μm^3^ for *in vivo 2p* and 0.154 ± 0.004 synapses/μm^3^ for *XTC Restored*, p < 0.001; unpaired two-tailed t-test). Moreover, we found that baseline synapse dynamics were quite stable across 10 days of imaging in *XTC Restored* volumes (Fig. 6f). However, the median fold change for non-restored volumes was not as stable, showing a marked decline on day 9 (Fig. 6b, f), suggesting that many tracking errors occurred on that timepoint. These errors are likely due to the accumulation of tissue shifts between the imaging session on day 2 and day 9 that were more difficult to resolve without XTC restoration. Moreover, when we manually assessed the rate of tracking errors for 100 randomly selected synapses before and after image enhancement, we also observed a reduced error rate at each individual timepoint, resulting in a 3-fold reduction in cumulative errors (Fig 6f, cumulative error of 18% for XTC and 61% for *in vivo 2p* on day 10). We also compared the distribution of sizes of all tracked synapses before and after image restoration and found that in both lateral and axial directions, the diameter of segmented synapses were much more similar to their expected physiological size (< 1 μm in diameter) in *XTC Restored* volumes as opposed to the raw *in vivo 2p* data (Fig. 6g - i; 1.0 ± 0.008 μm *Raw 2p* and 0.77 ± 0.005 μm *XTC* synapse equivalent diameter, 1.2 ± 0.009 μm *Raw 2p* and 0.77 ± 0.005 μm *XTC* max XY width, 1.0 ± 0.008 μm *Raw 2p* and 0.77 ± 0.005 μm *XTC* max XZ height, all comparisons p < 0.001; unpaired two-tailed t-test). Overall, this analysis indicates that image restoration facilitates the tracking of individual synapses across multiple days *in vivo* by de-noising and super-resolving images, simplifying the tracking task such that sparsely annotated simple tracking models can be effectively employed.

#### Cross modal image enhancement enables detection of behaviorally relevant synaptic changes

Accurate *in vivo* synapse tracking holds the strong potential to quantitatively relate changes in synapse strength within defined neuronal circuits to behavioral outcomes. To assess whether *XTC Restored* volumes can be used to track and detect a biological difference in an established behavioral paradigm for synaptic plasticity, we exposed mice to an enriched novel environment after 3 days of baseline imaging (Days 1-3) (Fig. 7a). Environmental enrichment (Day 4, occurring 2 hours before imaging session) consisted of a single, 5-minute period of exploration in a chamber containing novel visual, auditory, olfactory, and tactile stimuli, after which the mice were transferred back to their home cage. Animals were then imaged on days 5, 7, 9 and 11. To correct for shifts in global signal intensity, SEP intensity was normalized to the red signal of a sparse subset of neurons filled with tdTomato, as both signals were excited by the same beam and detected with high- and low-pass filters respectively (910nm 2p excitation; Fig. 7b).

**Figure 7.**
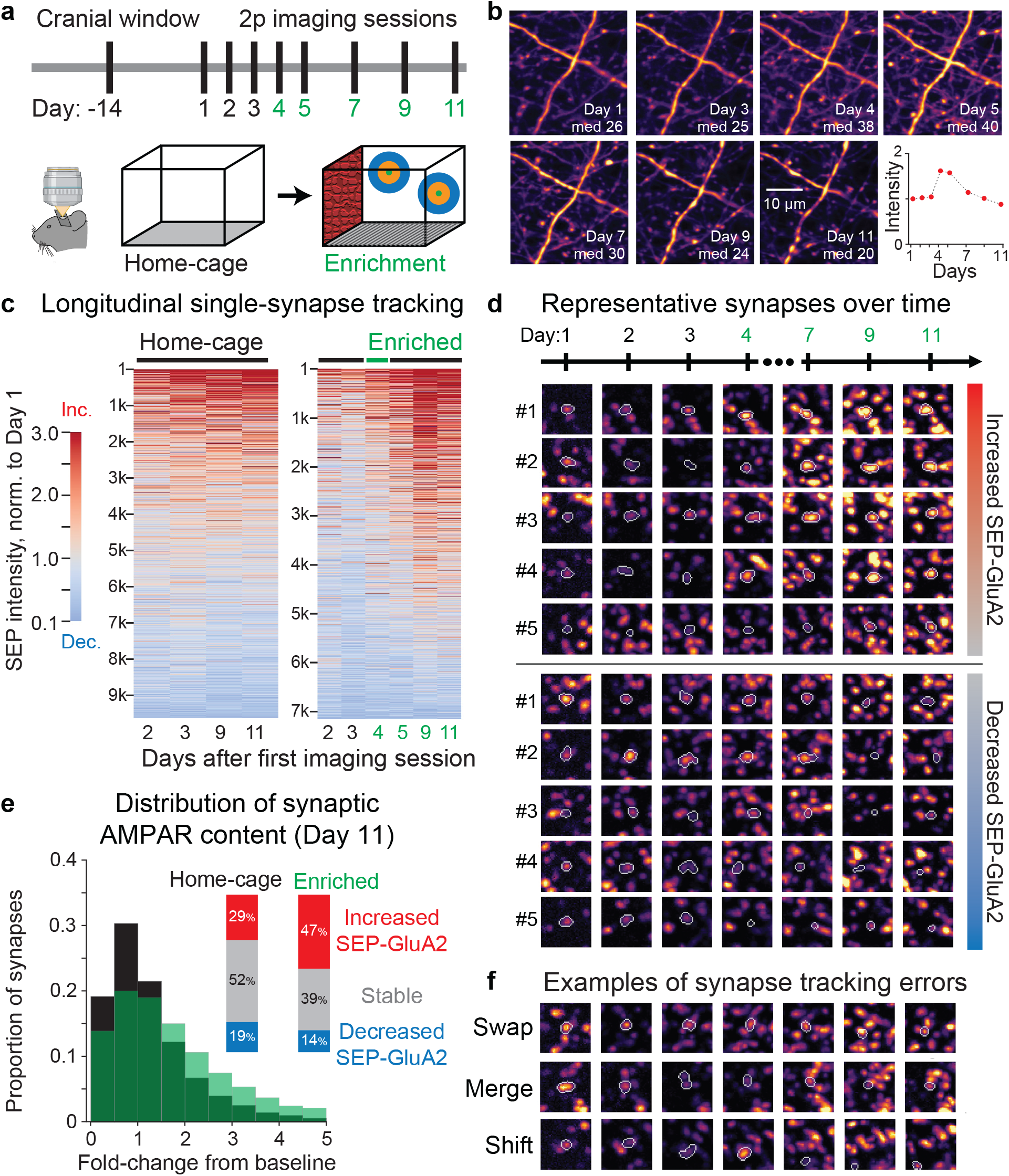
Increased synaptic plasticity in cortex following environmental enrichment. (**a**) Experimental timeline. Same volumes of retrosplenial cortex were imaged over 11 days. On Day 4, animals were exposed to enriched environment for 5 min, consisting of novel textures, smells, sounds, and visual cues. (**b**) Variations in daily signal intensity were measured and compensated by normalizing SEP signal to red cell fill, excited by the same beam path. (**c**) Comparison of synapse dynamics in home-cage (control) and enriched environment mice. Fold change relative to Day 1 baseline. 1 representative mouse displayed per condition. (**d**) Subset of tracked synapses that displayed increased or decreased strength relative to baseline. Maximum projection of cropped volumes (4×4×4μm). (**e**) Proportion of synapses that increased or decreased after 1 week of environmental enrichment. Inset, quantification of synapses that increased, decreased, or remained stable in control and experimental condition. Thresholds of >150% and <50% of Day 1 SEP intensity defined increased and decreased SEP-GluA2 expression on Day 11. (**f**) Examples error types in longitudinal synapse tracking.

Animals exposed to environmental enrichment showed a marked shift in the proportion of synapses with both stronger and weaker synaptic connections (corresponding to increased and decreased synaptic SEP-GluA2 content, respectively), and an overall net increase in synapse strength, consistent with the net induction of long-term potentiation (LTP), which is known to encode both spatial learning and environmental enrichment. Representative examples of successfully tracked synapses are shown in relation to the first imaging timepoint (Fig. 7d). The proportion of synapses that increased in strength, defined as having a sustained fold change > 1.5 on days 9 and 11, was 16.9% in home cage condition as compared to 24.3% in enrichment condition. Moreover, the proportion of synapses that decreased in strength, defined as having a sustained fold change < 0.5 on days 9 and 11, was also lower after enrichment (24.0% in home cage and 21.0% in enrichment, Fig. 7e). Notably, while home-cage control animals showed some synapse dynamics over time, rates of AMPAR addition and over 10 days was balanced, suggesting no net synaptic plasticity (Fig. 7c), and the magnitude of increased and decreased AMPAR content was significantly lower compared to animals that received environmental enrichment. The median fold-change and overall distribution of fold-changes for individual synapses (Fig. 7e) were all shifted to higher values after enrichment. However, while our simple *ilastik* tracking model was able to detect these biological differences, we still noted several errors that could be categorized into error-types that future models should focus on preventing: “swap” errors, where a tracked synapse incorrectly jumps to a nearby synapse at a later timepoint; “merge” errors, which occur at the detection stage, and result in blobs of synapses being tracked as a single entity; “shift” errors, where one association error will continue to propagate on additional timepoints in regions with poor registration (Fig. 7f). These can be improved with more complex detection and tracking models using deep learning or other machine learning approaches. Despite these tracking errors, *XTC Restored* volumes facilitated the tracking and detection of a behaviorally relevant change in synaptic strength *in vivo*.

## Discussion

Studying structure-function relationships is a classic paradigm to explore the fundamental basis of neural circuits. For example, connectomics seeks to generate comprehensive maps of entire nervous systems in order to establish the physical architecture that gives rise to neural function. However, these static wiring diagrams do not reveal dynamic processes that play important roles in learning and behavior, which are higher-order cognitive functions that are encoded largely as changes in synaptic strength in real-time. Thus, to fully understand how behavioral learning is encoded as changes in the strength of individual synapses within complex neural networks, it is crucial to develop tools that enable synapse tracking *in vivo*. While recent advances in molecular labeling have enabled visualization of the synaptome by fluorescently labeling structural components of the synapse (Zhu et al., 2018), these tools have only been applied *in vitro* to produce rich maps of synapse structure across the brain at a single static point in time. The dearth of tools that enable longitudinal tracking of synapses in living animals hinders our understanding of fundamental questions in neuroscience, such as elucidating the molecular mechanisms of information storage and behavior. By developing computational tools to enhance low-resolution 2p images to optimal Airyscan-level image quality, our study is the first to map the synaptome of large cortical regions *in vivo* and is also the first to explore the *functional connectome*, a comprehensive map of how synapses undergo plasticity during behavioral experience. Crucially, our transgenic labeling strategy tags the functional component of excitatory synapses, the AMPA receptor. Thus, our experimental paradigm can generate detailed maps of synaptic plasticity that are directly related to animal behavior, revealing how plasticity is encoded spatially and temporally in living animals. To enable the detection and tracking of synapses in these dense datasets *in vivo*, we developed a computational pipeline to register, superresolve, identify, and track millions of individual synapses automatically.

At the core of this novel paradigm for visualizing synapses is a cross-modality supervised image restoration model that offers several advantages. First, the XTC restoration model significantly improved the axial and lateral resolutions of *in vivo* 2p images, enabling Airyscan-like quality *in vivo* while retaining the benefits of increased depth and imaging speed associated with 2p excitation and resonance scanners. Moreover, the XTC restoration model was able to generalize across different data, such as fluorescently labelled neurons, validating the obtained network and demonstrating that it was not overfitting on SEP-GluA2 signals. Second, by enhancing the resolution of *in vivo 2p* images, the task of synapse detection and tracking was significantly simplified, such that established machine learning algorithms trained with sparse human annotations could reasonably segment and associate synapses relative to ground-truth *Slice Airy* data. In essence, this image restoration pipeline significantly denoises *in vivo 2p* data, thereby facilitating the identification and longitudinal tracking of synapses.

Beyond these important quantitative considerations, our cross-modality training and inference paradigm serves as a proof-of-concept for generating similar supervised restoration algorithms for the enhancement of other *in vivo* fluorescent signals. We showed that paired high-low resolution training data, acquired in live slice preparations, is sufficient to build a restoration model capable of superresolving signals *in vivo*. Using live slice tissue to generate training data is much more routinely feasible than having to acquire paired training data by registering the same field-of-view first *in vivo* and then post-dissection in slice, which is virtually impossible for nanoscale synaptic structures. Thus, using our slice training paradigm, researchers can rapidly generate trained CNNs to enhance a multitude of fluorescent signals *in vivo*, such as transient calcium signals, fluorescent molecules that are easily photo-bleached, or other nanoscale structures that are near the detection limit of *in vivo* objectives. Several emerging imaging applications in particular suffer from similar fundamental resolution barriers *in vivo*, including axon terminal imaging and new approaches to large-scale voltage imaging (Platisa et al., 2021). These advantages stem from the fact that XTC uses data from other modalities to train a signal model. Thus, XTC is a complementary approach to other neural network designs that focus on using noise statistics for denoising imaging data (Lecoq et al., 2021).

A final major advantage of this CNN-based restoration model is its ability to adapt to local variations in image statistics without relying on heavy data pre-processing. Traditional contrast enhancement algorithms, such as CLAHE, tend to work best when the overall illumination in an image is uniform (Pizer et al., 1990). Otherwise, bright signals will become over-saturated if the global image is dim, and faint signals will remain dim if the global image is bright. In our case, supervised image restoration adapted to the local image features in dim, dense, and bright regions automatically, thereby preserving the overall distribution of intensity values without over-saturating any voxels. No additional pre-processing was necessary when applying the trained model to new data beyond image interpolation to ensure that the image resolution was within the working range.

A caveat of this supervised restoration paradigm is that it is fundamentally limited—as expected— by the resolution of optical elements. For instance, most synapses that were not detected in *XTC Restored* volumes were below the diffraction limit (Fig. 5g, h). Thus, XTC processing is simply improving the signal-to-noise ratio of signals that already exist within our *in vivo* 2p data. Moving beyond the diffraction limit requires either better objectives and optical components that can acquire more signal (Steffens et al., 2020; Wegner et al., 2018) or computational imaging techniques that use coded apertures and algorithmic signal reconstruction (Aidukas et al., 2019; Kazemipour et al., 2019; Song et al., 2017). Furthermore, additional improvements could be realized by expanding on the synapse detection and tracking algorithms currently trained in *ilastik*. In the present XTC pipeline, employing simple classifiers minimized manual labor, increased algorithmic availability, and enabled us to showcase the extent to which the core image restoration algorithm facilitates synapse segmentation and tracking. We expect that as synaptic imaging progresses, more sophisticated algorithms will be explored. For instance, 3D Mask-RCNN models (He et al., 2017) perform exceptionally well at segmenting individual objects in regions of high-density, thereby circumventing the need for watershed segmentation after voxel classification. Moreover, for synapse tracking, the application of sophisticated registration algorithms, like LDDMM (Beg et al., 2005), can further trivialize the challenge of tracking. Other algorithms, like *DeepSort* (Wojke et al., 2017), can be trained to resolve any tracking errors that persist. A final limitation that often occurs during synapse tracking is the slight shifting of large blood vessels across timepoints. While these shifts do not impact registration, they do obscure any synapses that lie underneath the vessel, thus artificially reducing the intensity of detected SEP synapses. Currently, these synapses are not included in our analysis, but future improvements can introduce static landmarks, such as injected fluorescent beads, to better account and normalize for local shifts in intensity, allowing researchers to recover obstructed signals.

Overall, the biological and computational tools presented here enable fundamentally new experimental paradigms. By training and validating CNNs using multi-modal imaging, we provide a framework to restore optimal Airyscan-like resolution to *in vivo* 2p images that provide new insight into synapse dynamics encoding behavioral learning over extended time scales. Our proof-of-concept experiments illustrate that these tools can be used to track behaviorally relevant plasticity with single-synapse resolution at a brain-wide scale. More generally, these tools can be applied to investigate synaptic plasticity in any brain region during any behavioral paradigm. Beyond these important observations of the synaptic basis of behavior, our work makes accessible the richness of *in vivo* longitudinal datasets generated from SEP-GluA2 transgenic animals. Using XTC, future experiments can now more easily advance detection and tracking algorithms that will enable both temporal and spatial exploration of synaptic plasticity. For instance, detailed spatial analysis can identify the location of subpopulations of synapses that change their synaptic strength in concert across cortical regions. Moreover, combined with Cre-dependent neuronal labeling, researchers can study synaptic plasticity within specific subsets of neurons in the brain. Thus, our computational cross-modality image restoration paradigm sets the stage for detailed molecular studies of synaptic plasticity underlying behavior, bringing us a step closer to understanding the foundations of cognition.

## Methods

### Mouse genetics

SEP-GluA2 mice were made in collaboration with the Johns Hopkins University School of Medicine Transgenics core. Briefly, we used a CRISPR-Cas9-based approach to insert a SEP tag into exon 1 of *Gria2*, localized to the area encoding the GluA2 subunit N-terminus. Homozygous transgenic mice are viable, breed well, and appear to be physiologically and behaviorally normal. In this transgenic line, all GluA2-containing AMPARs are labeled at endogenous levels, enabling robust visualization of excitatory synapses throughout the entire brain.

### Surgical procedures

Mice were anesthetized (1.5-2% isoflurane) and implanted with a 3×3 mm cranial window (Potomac Photonics) over retrosplenial cortex at 10-16 weeks of age. Windows were sealed and custom-made metal head bars attached using dental cement (Metabond; Edgewood, NY). In a subset of experiments, an AAV-CaMKII-cre virus (Addgene/Penn Vector) was injected into cortex (viral titer: 5e8-1e9, 100-150 nL, 0.3 mm deep) of double homozygous SEP-GluA2 x Ai9 reporter mice to sparsely label L2/3 pyramidal neurons with a tdTomato cell fill. 10 mg/kg of extended-release Buprenorphine (ZooPharm) was administered before surgery and mice were observed for 3 days following surgery. Mice were allowed to recover for at least 2 weeks before commencing *in vivo* imaging. All surgical procedures were approved by the Johns Hopkins Animal Care and Use Committee.

### Mouse behavior

Mice were handled daily for 1 week prior to behavioral testing. Environmental enrichment consisted of a single 5-min exploration session in a novel spatial environment. Mice were individually placed in a 20cm square chamber with distinct spatial markings and textures on the walls (60-grit sandpaper), a novel smell (70% acetic acid), novel cage floor (1-cm-spaced circular bars), and white noise (70dB). Home-cage control mice were handled daily for one week before the start of the experiment but were not exposed to novel spatial environment.

### In vivo and in vitro 2p imaging

*In vivo* 2p images were acquired from lightly anesthetized mice (1.5% isoflurane) using a custom-built, 2p laser-scanning microscope controlled by ScanImage (Vidrio, Ashburn, VA) and a 20×/1.0 NA water-immersion objective lens (Zeiss). SEP-GluA2 (green) and tdTomato cell fill (red) were both excited at 910 nm with a Ti:sapphire laser (SpectraPhysics, 20 mW power at objective back-aperture). Green and red fluorescence signals were acquired simultaneously and separated by a set of dichroic mirrors and filters (ET525/50m for green channel, ET605/70m for red channel, Chroma). Image stacks were acquired by resonance scanning at 30-Hz. The field-of-view contained 1,024 × 1,024 × 70 voxels with a lateral XY resolution of 0.096 μm/px and an axial resolution of 1 μm/px. Live, 300 μm thick acute slices of SEP-GluA2 brains were imaged using the same optical setup, except that the edges of the tissue were embedded on a microscope slide with 4% w/v gelatin, as described for single-photon imaging. Slices were maintained in HEPES-buffered ACSF, consisting of: 140 mM NaCl, 5 mM Kcl, 10 mM glucose, 10 mM HEPES, 2 mM CaCl2, 1 mM MgCl2, pH adjusted to 7.40.

### Single-photon confocal and Airyscan imaging

Paired high-resolution Airyscan and low-resolution confocal training volumes were generated using a Zeiss 880 microscope and a 63×/1.0 NA water objective lens (Zeiss) in live slice preparations. Live, 300 μm thick acute slices of SEP-GluA2 brains were embedded to microscope slides by applying a warm 4% w/v gelatin solution along the edges of the tissue and then cooling the gelatin at 4°C until solid. This embedding prevented the tissue from drifting during ACSF-immersion imaging and helped preserve the endogenous SEP-GluA2 signal. SEP-GluA2 and tdTomato cell fill were excited at 488 nm and 546 nm, respectively. Optimal high-resolution images were acquired using calibrated Airyscan detectors that achieved a lateral resolution of 0.063 μm/px and an axial resolution of 0.33 μm/px. Immediately after high-resolution volumes were imaged, paired suboptimal images were acquired to reduce registration errors. The image quality of suboptimal images was curated to replicate the image quality of *in vivo* 2p datasets by opening the confocal pinhole 4 times higher than the ideal A.U., increasing the laser gain to near maximal levels, and reducing laser power. Overall, we collected 24 paired high-low resolution training volumes with a 550 × 550 × 20 voxel field-of-view from 8 tissue slices containing multiple cortical regions. Validation and all *in vivo* data were generated from different animals (n=5 mice). In addition to live-slice imaging, we also applied the Zeiss 880 microscope in Airyscan Fast mode (Huff, 2016), with a 20×/1.0 NA water-immersion objective lens (Zeiss), to attempt to detect synapses *in vivo*.

### Neural network architecture and optimization

Having collected pairs of aligned high- and low-resolution images, 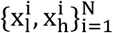 we employed a supervised learning approach in order to find a map from input images x_1_ ∈ R^n_l_^ to outputs x_h_ ∈ R^n_h_^, where n_l_ < n_h_. In particular, we parametrize this function *f*_θ_: *R^n_l_^* → *R^n_h_^* with a convolutional neural network with parameters θ. We employ a convolutional neural network architecture similar to those proposed by Weigert et al with a modified U-Net architecture (Ronneberger et al., 2015; Weigert et al., 2018). Following (Dong et al., 2015), the input is first bicubic-interpolated to match the target dimension before applying a function parameterized by the CNN. At deployment, *in vivo* volumes follow an analogous pipeline whereby they are first interpolated to match the axial and lateral resolutions of the training data before restoration. Following an empirical risk minimization approach, we minimize a suitable loss function over the training samples according to:

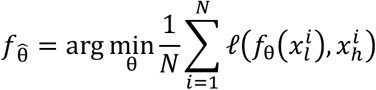

The loss penalizes deviations by the reconstructed images, 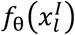, from the high-quality samples, 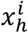. For simplicity, we chose to optimize the average of mean-absolute error (MAE), namely 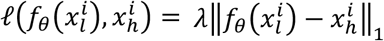. Combining this with other losses, such as the multi-scale structural similarity index (MS-SSIM) (Wang et al., 2003), is certainly possible and might provide further improvements. Overall, we trained a convolutional neural network with paired high-resolution *Slice Airy* and low-resolution *Slice 1p* data for 1000 epochs with batch size of 8 on a NVIDIA Tesla P100-PCIE GPU using Adam optimizer (Kingma and Ba, 2014) and a learning rate of 4 × 10^-4^.

### Validation comparisons

The ideal validation experiment would require imaging the same field-of-view first *in vivo*, using our 2p setup, and then again *ex vivo* in slice, with Airyscan detectors. This setup would allow us to directly compare synapses detected in *XTC Restored in vivo* images to synapses in ground-truth *Slice Airy* volumes. Unfortunately, we found that it was extremely difficult to find the exact same field-of-view *ex vivo* post-perfusion, even with the addition of structural anchors to help with registration, such as sparse neuronal labels. Moreover, we found that it was entirely impossible to preserve the tissue throughout dissection such that the position of synapses remained stable enough for registration. Thus, we chose to perform our validation in live slice tissue directly, where we could try to faithfully replicate the quality of images acquired *in vivo* by using the same 2p microscope, resonance scanner, objective, and laser power to acquire our low-resolution *Slice 2p* images (Fig. 2b,c). Moreover, tissue slices were imaged immediately after dissection to preserve endogenous tissue quality. The high-resolution images (*Slice Airy*) paired to these low-resolution *Slice 2p* volumes were acquired on a separate Zeiss 880 microscope equipped with Airyscan detectors (Fig. 2). The high- and low-resolution validation pairs were then registered together using a combination of FIJI’s correct 3D drift package (Parslow et al., 2014) and *SimpleElastix* affine transformations (Marstal et al., 2016) to remove tissue movements that occurred when transferring between microscopes.

Finally, we compared pairwise detected synapses, segmented using *ilastik* voxel classifiers, between high-resolution *Slice Airy* volumes and both *Slice 2p* and *XTC Restored* volumes. For these comparisons, we defined “true positive” detections as synapses sharing at least one voxel across the pairwise compared volumes. As this threshold was very lenient, we also included several validation metrics to assess pairwise structural and intensity similarities to validate the extent to which XTC processing improved the size and shape of synaptic detections.

### Registration and data processing for synapse detection and longitudinal tracking

Several pre-processing steps were applied to facilitate longitudinal synapse tracking (Fig. 2d). First, each image at a given timepoint *t* was volumetrically registered to the subsequent timepoint *t* + 1 using affine transformations in *SimpleElastix*. While volumetric registration accounted for global tissue shifts, we found that local misalignments persisted after volumetric registration. Thus, we included an additional registration step that registered each XY slice on timepoint *t* with each corresponding slice on timepoint *t + 1*. The final pre-processing step was to detect blood vessels and exclude them from our analysis, as synapses located adjacent to blood vessels could easily become obscured and appear as an eliminated synapse. To perform blood vessel masking, volumes were binarized, followed by inversion, image opening and dilation to extract a smooth binary mask that excluded dim dark regions. No other pre-processing steps were applied to the raw data, and the registered volumes were then processed with XTC.

Synapse detection and tracking algorithms were trained in *ilastik*, a platform that enables researchers to rapidly build machine-learning models using sparse annotations. For synapse detection, random-forest classifiers were trained, each using 30 human segmented synapses, for all imaging modalities (*Slice 2p, Slice Airy, in vivo 2p, XTC Restored*) respectively. For synapse tracking, we trained two models using structured sparse learning in *ilastik*, each with 100 human annotated tracks, for *in vivo 2p* data both before and after XTC restoration. To ensure that the intensity of the SEP signal compared across tracking experiments is not altered by XTC processing, we also overlaid the XTC segmentations on the *in vivo 2p* data to extract intensity values directly from the raw data.

### Statistical Analysis

All statistical analysis was performed using Python statsmodels and scipy libraries. N represents the number of animals used in each experiment, unless otherwise noted. Data are reported as mean ± SEM or median ± SEM as indicated, and *p* < 0.05 was considered statistically significant. Level of significance is marked on figures as follows: * denotes *p* < 0.05; ** denotes *p* < 0.01; *** denotes *p* < 0.001.

### Code Availability

Packaged software code will be made available along with instructions for use and demo data across a small volume. The algorithm is prepared to work with minimum Python 3.6.

## Acknowledgements

We would like to thank Michael Miller and Daniel Tward for their work on an earlier version of the automatic synapse detection pipeline. We would also like to thank Lucien E. Weiss for constructive comments on an earlier version of this work. YKX and ARG are supported by Kavli Neuroscience Discovery Fellowships. ARG and RLH are supported by R21 AG063193 and R01 MH123212. Additional funding was provided by R01 RF1MH121539.

---

**Video 1**. **Cross-modality image enhancement improves synapse resolution *in vivo***. Identical volumes of SEP-GluA2 tissue imaged *in vivo*, before (left) and after (right) applying XTC. 70μm depth, 1μm steps, starting from pial surface and moving deeper into brain.

## Notes

### Competing Interest Statement

The authors have declared no competing interest.

